# Two opposing redox signals mediated by 2-Cys peroxiredoxin shape the redox proteome during photosynthetic induction

**DOI:** 10.1101/2025.04.02.646757

**Authors:** Shani Doron, Nardy Lampl, Alon Savidor, Amir Pri-Or, Corine Katina, Francisco Javier Cejudo, Yishai Levin, Shilo Rosenwasser

## Abstract

Photosynthetic induction, characterized by the lag in CO_2_ assimilation rates typically observed upon plant transition from darkness to light, has traditionally been attributed to Rubisco activase activity and stomatal opening. Yet, the faster induction of photosynthesis in the 2-Cys peroxiredoxins (Prxs) mutant (*2cpab*) highlighted the critical role of chloroplast redox state in regulating photosynthetic rates during this phase. Since 2-Cys Prxs are involved in transmission of oxidative signals to target enzymes, it was hypothesized that it slows down photosynthesis during the induction phase. SPEAR, a redox proteomics approach for simultaneous protein expression and redox analysis, was used to systematically map redox changes occurring at the proteome level during photosynthesis induction and to unravel the role of 2-Cys Prxs in shaping these redox alterations. No significant difference was observed in protein expression levels between WT and *2cpab* plants, suggesting that protein abundance does not account for the *2cpab* phenotype. During the transition from dark to low light, 82 and 54 cysteine-containing peptides were reduced or oxidized, respectively, in WT plants. Most redox-regulated cysteines in photosynthetic proteins were found oxidized in the dark and became reduced in response to light, including ATP synthase gamma chain 1 (ATPC1) and glyceraldehyde-3-phosphate dehydrogenase (GAPB). A reverse pattern was observed among redox-regulated cysteines in proteins involved in starch degradation and chloroplast glycolysis, which shifted from a reduced to an oxidized state in response to light. These findings demonstrate the initiation of two opposing redox responses, affecting distinct sets of metabolic proteins during the induction phase. Remarkably, a significantly lower number of cysteines were reduced or oxidized in *2cpab* plants, highlighting the crucial role 2-Cys Prxs play in shaping both signals. Taken together, rotational shifts between metabolic pathways during the photosynthesis induction phase are regulated by two opposing redox signals mediated by 2-Cys Prx activity.

## Introduction

Converting light into chemical energy through the light and carbon reactions of photosynthesis is a highly intricate process, demanding the orchestration of numerous enzymatic reactions. This coordination is especially critical under the dynamic light conditions experienced by plants, which cycle between full sunlight, shade and complete darkness over the course of a day, driving continuous activation and deactivation of the photosynthetic process. The importance of coordinating reaction rates under dynamic light conditions arises from the high vulnerability of the photosynthetic electron transport chain to over-reduction and reactive oxygen species (ROS) production when the absorption of light energy, and the resulting production of NADPH and ATP, exceed the capacity of downstream reactions, necessitating the activation of energy dissipation mechanisms and antioxidant defenses^1–3^.

Redox modifications driving the reconfiguration of chloroplastic metabolic activities in response to changes in photosynthetic performance evolved in plants to maintain homeostasis and prevent the accumulation of harmful ROS. Specifically, thiol-based regulation allows communication between the photosynthetic light reaction and downstream metabolic activity by transferring redox signals that finetune the activity of various enzymes^4–9^. Activation of redox-regulated enzymes is dependent on reductive signals that drive disulfide bond reduction and are transmitted from ferredoxin to thioredoxins (Trxs) via ferredoxin-Trx reductase (FTR) and subsequently from Trxs to target proteins^5,10–12^. Reductive signals are also transmitted through ferredoxin-NADP(+) reductase (FNR) and NADPH Trx reductase C (NTRC)^13–15^, broadening the range of redox-regulated proteins and the sources for reducing power. Counterbalanced to the reductive activity, disulfide bond formation depends on the transfer of reducing equivalents from target proteins to H_2_O_2_, catalyzed by high midpoint potential atypical Trxs and 2-Cys-Prxs^16–21^. While the reductive activity catalyzing the reduction of disulfide bonds is typically associated with light, oxidative signals have mainly been studied in the context of light-to-dark transition, where they catalyze the formation of disulfide bonds, resulting in enzyme deactivation^8,9^.

The induction phase of photosynthesis is characterized by a lag in the rise of photosynthesis to the maximum light-saturated rate, which typically occurs during dark-light or low-high light transitions. The relatively slow kinetics of photosynthetic activation, which requires several minutes for readjustment, has been suggested to cause a substantial loss of potential photosynthetic capacity in plants experiencing dynamic light conditions^22,23^. Multiple factors influence the speed of induction, including activation of Rubisco through Rubisco activase, and stomatal opening, with the fast phase of the induction process controlled by the rate of enzyme activation by the Fd/Trx system and optimization of the electron transport chain^24–28^. Characterization of the reductive activation of various chloroplast enzymes during dark-to-light transitions^29,30^ has highlighted the role of thiol-based regulation in shaping the kinetics of the photosynthetic induction process. Interestingly, significantly faster induction of photosynthesis during the transition of plants from dark to low light (LL) was measured in Arabidopsis (*Arabidopsis thaliana*) plants lacking 2-Cys Prxs A and B (*2cpab*) in comparison to wild type (WT)^31^, pointing to the inhibitory effect of these enzymes during the photosynthesis induction phase. However, the mechanism by which 2-Cys Prxs suppress photosynthetic activity remains unclear.

By characterizing redox changes occurring at the proteome level during the photosynthesis induction phase, this work identified 136 Cys sites that underwent rapid redox changes in response to a minute amount of light in WT plants. Many of the redox changes observed in WT plants were absent in *2cpab* plants, marking 2-Cys Prx as a key regulator of protein dynamics during the initiation of photosynthesis. Notably, the transition from dark to light simultaneously triggered two opposing 2-Cys Prx-regulated redox responses, with some target proteins undergoing reduction and others undergoing oxidation. While most Calvin–Benson cycle (CBC) and light reaction enzymes were reduced in response to light, downstream metabolic pathways, such as starch degradation, were oxidized. These results suggest that dark-to-light transitions involve dynamic shifts in metabolic pathways, governed by opposing redox signals.

## Results

### Global quantification of redox-sensitive cysteines during the photosynthesis induction phase

The SPEAR labeling approach, which allows for simultaneous protein expression and redox analysis,^32^ was used to map proteome-level redox changes occurring during the photosynthesis induction phase and elucidate the role of 2-Cys Prx in shaping oxidation patterns during this phase (Fig. 1a). SPEAR is based on the rapid acidification of plant extracts with trichloroacetic acid (TCA) to preserve the redox status of thiol groups, followed by differential labeling of reduced and reversibly oxidized cysteines with isotopically light (d_0_) and heavy (d_5_) forms of N-ethylmaleimide (NEM), respectively. Quantification of the relative abundance of the reduced and oxidized forms is achieved by comparing the area under the curve of the light- and heavy-labeled peptides at the intact peptide level (MS1), enabling the calculation of the oxidation degree (OxD) for each identified cysteine site. As low-abundance oxidized peptides (d_5_-NEM) might limit the number of identified peptides, an additional set of samples in which the reduced Cys groups are labeled with d_5_-NEM is analyzed in parallel to the experimental samples, serving as markers to locate m/z and retention time (RT) values of low-abundant, reversibly oxidized peptides. As no enrichment of cysteine-containing peptides is included, data regarding protein expression levels can also be extracted from the MS spectra.

**Figure 1:**
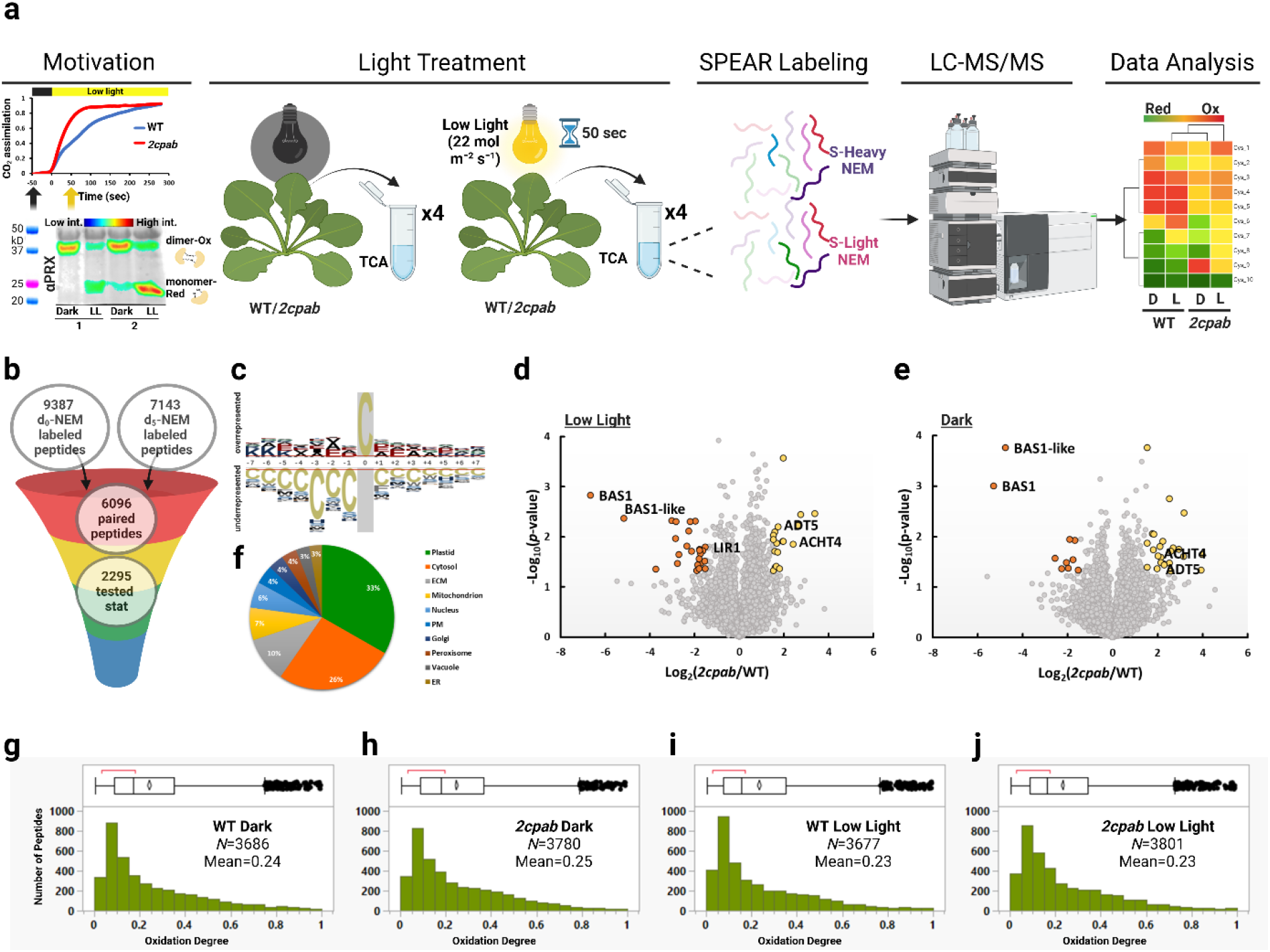
Simultaneous quantification of Cys oxidation and protein expression in WT and *2cpab* plants under dark and low light conditions. **(a)** Schematic representation of the workflow for this experiment. The upper graph in the Motivation part, was adapted from Lampl et al. (2022). **(b)** The number of identified d_0_-NEM-labeled peptides, d_5_-NEM-labeled peptides, peptides labeled with both d_0_ and d_5_-NEM and paired peptides detected in at least two replicates across all treatments. **(c)** The consensus motif of NEM-labeled peptides. The *Arabidopsis thaliana* proteome was used as a background population for probabilities calculation. A similar analysis performed for non-labeled Cys-containing peptides is presented in Supplemental Fig. 1. (**d, e)** Volcano plot visualization of the expression profile of 9662 proteins in *2cpab* compared to WT plants following treatment with low light **(d)** or dark **(e)**. Colored dots highlight proteins with an average ≥ 1.5 log2-fold change in *2cpab* mutant plants compared to WT plants and a p-value < 0.05. **(f)** Subcellular distribution of the identified thiol proteins was determined using the SUBA tool. **(g-j)** Distribution plots displaying the oxidation degree of the peptides found in **(g)** WT dark, **(h)** *2cpab* dark, **(i)** WT low light and **(j)** *2cpab* low light plants.

Samples were taken from WT and *2cpab* plants, both kept in the dark and following a transition to LL conditions (22 μmol m^-2^ s^-2^), simulating the transition from night to the beginning of daylight. Furthermore, sampling under LL, when the redox carriers of photosynthetic reactions are not saturated, minimizes the influence of other potential limiting factors on CO_2_ assimilation rates^29^. To verify the sensitivity of 2-Cys Prx to LL, its oxidation state was measured by calculating the relative amounts of the dimeric (oxidized) versus monomeric (reduced) forms on non-reducing SDS-PAGE using a 2-Cys Prx-specific antibody. As shown in Fig. 1a, 2-Cys Prx switched from its oxidized state to its reduced state after exposure to LL, verifying its responsiveness to relatively low electron flux.

To capture the early molecular events that might shape the gap between WT and *2cpab* photosynthesis rates, samples were taken 50 sec following the transition to LL, 20 sec before the difference in CO_2_ assimilation rates between the two lines reached its maximum^31^ (Fig. 1a). In total, 9387 d_0_-NEM and 7143 d_5_-NEM-labeled peptides were identified, 6096 of which were labeled with both NEM isotopes, representing proteins from various subcellular localizations (Fig. 1b). Notably, most (∼96%) of the identified Cys-containing peptides were found to be NEM-labeled, reflecting the high efficiency of the labeling procedure. In agreement with our previous observation^32^, the NEM-labeled cysteine residues were found in close proximity to a consensus motif significantly enriched with glutamic acid and aspartic acid (Fig. 1c). This motif was absent in Cys-containing peptides that remained unlabeled by NEM (Supplemental Fig. 1). These observations suggest that glutamic acid and aspartic acid residues enhance cysteine reactivity to NEM and align with their suggested role in enhancing Cys nucleophilicity by modulating its pKa^33,34^.

Examination of the difference in protein expression in WT vs. *2cpab* plants under either LL or dark conditions (Fig. 1d and 1e, respectively), identified altered expression levels of only a few proteins in the *2cpab* compared to WT (|Log2 FC|>1.5, p-value < 0.05, Supp. Table 1). Among the chloroplastic proteins, there was an upregulation of the atypical cysteine histidine-rich Trx 4 (ACHT4) protein in *2cpab*. As ACHT4 oxidation is driven by 2-Cys Prx^18^, upregulation of ACHT4 may represent a mechanism aimed at compensating for the absence in 2-Cys Prx. In addition, Arogenate dehydratase 5 (ADT5) and light-regulated protein 1 (LIR1) were upregulated in *2cpab* plants (Fig. 1d, e). Overall, the slight difference in protein abundance between WT and *2capb* suggests that variations in photosynthetic efficiency between the lines are not regulated at the transcriptional or translational level, but rather, through post-translational modifications. Moreover, these results indicate that the mutation in 2-Cys Prx does not lead to H_2_O_2_ overproduction, as ROS accumulation is typically associated with a distinct transcriptional signature that should be reflected in protein expression^35–37^.

### 2-Cys Prxs shape the redox proteome during the photosynthesis induction phase

Only peptides detected in at least two out of four WT or *2cpab* plants replicates per condition (dark and LL) were used for the redox analysis, resulting in 2295 peptides, representing 1571 distinct proteins. Subcellular localization of the proteins, predicted using the SUBA tool^38^, was primarily in the chloroplast (33%) and cytosol (26%) (Fig. 1f). The mean oxidation degree (OxD) of cysteines in WT-dark, *2cpab-*dark, WT-LL, and *2cpab*-LL plants was 24%, 25%, 23% and 23%, respectively, demonstrating that the majority of the Cys residues remained highly reduced, irrespective of light conditions and genetic background (Fig. 1g-j).

Analysis of the oxidation degree difference (ΔOxD) between dark and LL conditions (OxD dark-OxD LL) in WT plants revealed 136 cysteine-containing peptides, representing 129 proteins that exhibited a significant change in their oxidation degree (|ΔOxD|>5% and P-value < 0.05) during the transition to LL in WT samples (Fig. 2a and Supp.Table 2). Functional annotations of the identified redox-regulated proteins indicated the enrichment of chloroplastic and stromal proteins (GO:0009507, GO:0009570), and those involved in photosynthesis (GO:0019253). Among the Cys sites that were redox modified in response to dark-LL transition, 18 were also found to undergo redox modifications in response to exogenous application of H_2_O_2_^32^, reinforcing their sensitivity to redox signals (Supp.Table 2).

**Figure 2:**
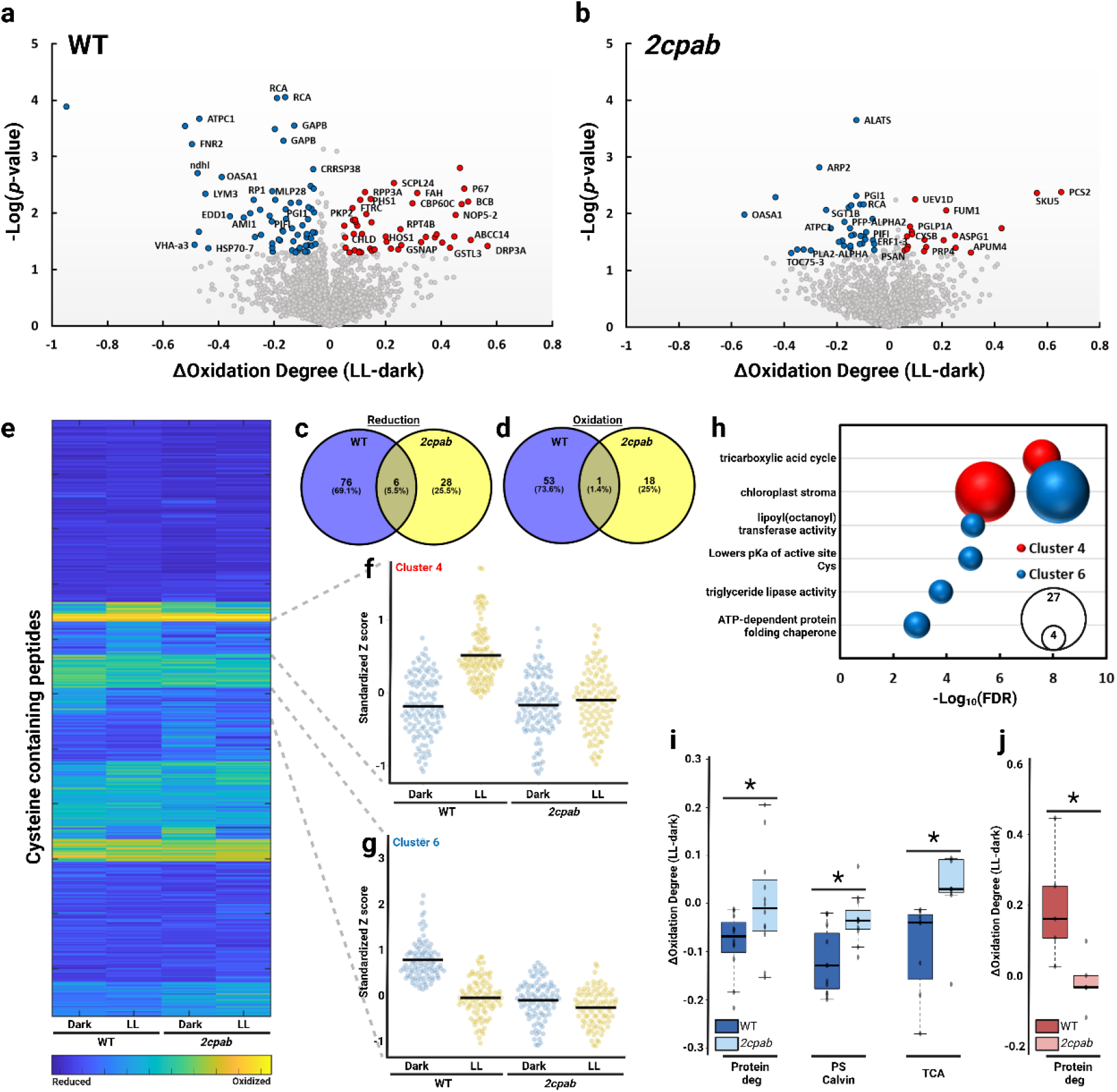
Photosynthetic induction is characterized by two opposing redox signals, both mediated by 2 Cys Prx. (a, b) Volcano plot visualization of the response of unique cysteine-containing peptides to the transition from dark to low light (LL) in the WT (a) and *2cpab* (b) plants. Colored dots highlight Cys residues with a ≥ 5% change in oxidation (ΔOxD) and a p-value < 0.05 in LL-treated plants compared to plants in the dark. (c, d) Venn diagrams depicting the reduced (c) and oxidized (d) unique cysteine-containing peptides in the transition from dark to LL in WT and *2cpab* plants. (e) Heat map visualization of K-means clustering of Cys oxidation state in response to LL in WT or *2cpab* plants. (f, g) Violin plots of the two highlighted clusters taken from the K-means clustering displayed in (e). (h) Significantly enriched GO terms and KEGG pathways (hypergeometric test, p < 0.05) in the clusters displayed in (f, g). Bubble size indicates the number of proteins that are associated with the indicated biological function. (i, j) Box plots showing changes in the redox state of reactive cysteines that underwent reduction (i) or oxidation (j), grouped by selected biochemical pathways, in WT and *2cpab* plants.

Among the redox-regulated Cys sites, 82, showed a significant reduction in response to LL, reflecting the transmission of reducing equivalents that activate proteins (Fig. 2a, c). Among them are Cys^241^ and Cys^248^ (ΔOxD = -13%, -11%, respectively) of both forms of 2-Cys Prx, A and B, and various Cys sites in photosynthetic proteins (Supp. Table 2, see more details later). Interestingly, redox modifications in response to LL were not restricted to reduction, as 54 cysteine-containing peptides exhibited oxidation upon exposure to LL (Fig. 2a, d). The identification of two distinct groups of cysteines with opposing responses suggests the concurrent transmission of both reducing and oxidative signals during the induction phase.

Significantly fewer redox modifications were observed during the induction phase in *2cpab* plants, with only 53 cysteine-containing peptides, corresponding to 53 proteins, showing a significant change in their oxidation state (Fig. 2b-d and Supp. Table S2). Notably, a smaller number of redox-modified Cys sites were identified in *2cpab* in both the reduced and oxidized groups, highlighting the crucial role of 2-Cys Prxs in mediating reductive and oxidative signals.

K-means clustering revealed two key patterns that demonstrated the role of 2-Cys Prxs in shaping the redox proteome during the induction phase (Fig. 2e). The first trend, reflected in the redox changes found in clusters 2 and 4, which were enriched with chloroplast stroma protein (GO:0009570), demonstrated reactive Cys sites that underwent oxidation in response to LL treatment in WT plants but remained reduced in *2cpab* plants, similar to their state in darkness (Fig. 2f, h and Supp. Fig. 3). This suggests a crucial role for 2-Cys Prxs in facilitating oxidation under light conditions. The second trend, reflected in the redox changes found in clusters 5 and 6, which were also enriched with chloroplast stroma proteins (GO:0009570), demonstrated reactive Cys sites that underwent reduction in response to LL treatment in the WT. In the *2cpab*, those reactive Cys were found to be already reduced in the dark and they also remained reduced in the light (Fig. 2g, h and Supp. Fig. 3). These data demonstrate that the lower number of Cys sites undergoing reduction in *2cpab* plants results from their lack of oxidation in the dark, aligning with the role of 2-Cys Prxs in protein oxidation during darkness^19–21^. Interestingly, clusters 8 and 10 comprised Cys sites that exhibited similar reduction and oxidation responses in both WT and *2cpab* plants (Supp. Fig. 3). This suggests that their redox regulation is unaffected by 2-Cys Prx, implying the presence of an alternative pathway for protein oxidation that operates independently of 2-Cys Prxs.

Mapping redox-regulated proteins to metabolic pathways and analyzing pathway-specific ΔOxD values further highlighted the role of 2-Cys Prxs during the induction phase. For example, redox-regulated peptides associated with CBC, which underwent reduction in response to LL in WT plants, exhibited a weaker reduction response in *2cpab* plants (Fig. 2i). Conversely, redox-regulated cysteines in proteins involved in protein degradation, which were oxidized in WT plants under LL, remained unoxidized in *2cpab* plants (Fig. 2j). The findings quantitatively demonstrate the role of 2-Cys Prx in mediating both reductive and oxidative signals across distinct metabolic pathways.

### Redox modifications in photosynthetic enzymes

Several cysteines in proteins involved in photosynthetic reactions were oxidized in the dark and became reduced in LL in WT plants, but were reduced in both the dark and LL conditions in *2cpab* plants (Fig. 3 & 4). This pattern indicates Cys oxidation in the dark, which is entirely dependent on 2-Cys Prx activity. Examples of cysteines that followed this trend in light reactions are Cys^141^ of Chlorophyll a-b binding protein CP26 (LHCB5), Cys^48/51^ of Photosystem I iron-sulfur center (psaC) and Cys^74^ of NAD(P)H-quinone oxidoreductase subunit I (ndhI) which are part of the iron sulfur cluster binding sites. Of note, the active site of TrxL2.2 and TrxL2.1, was reduced (Cys^113^/Cys^133^, ΔOxD = -30%) during the transfer from dark to LL in WT plants. In *2cpab*, compared to WT, this Cys was highly reduced in the dark and remained reduced when transferred to LL (Fig. 3). The lack of oxidation in *2cpab* plants in the dark aligns with the recent finding that TrxL2 is involved in the oxidation of chloroplast proteins in the dark by transferring reducing equivalents from target proteins to H_2_O_2_ through the activity of 2-Cys Prxs^20,39^. Moreover, considering the target proteins as the source of the reducing power for TrxL2, its reduction under LL suggests that, alongside electron transfer from photosynthesis electron transport chain to target proteins via the reductive pathway, a counterbalancing oxidative pathway simultaneously directs reducing equivalents away from target proteins to H_2_O_2_.

**Figure 3:**
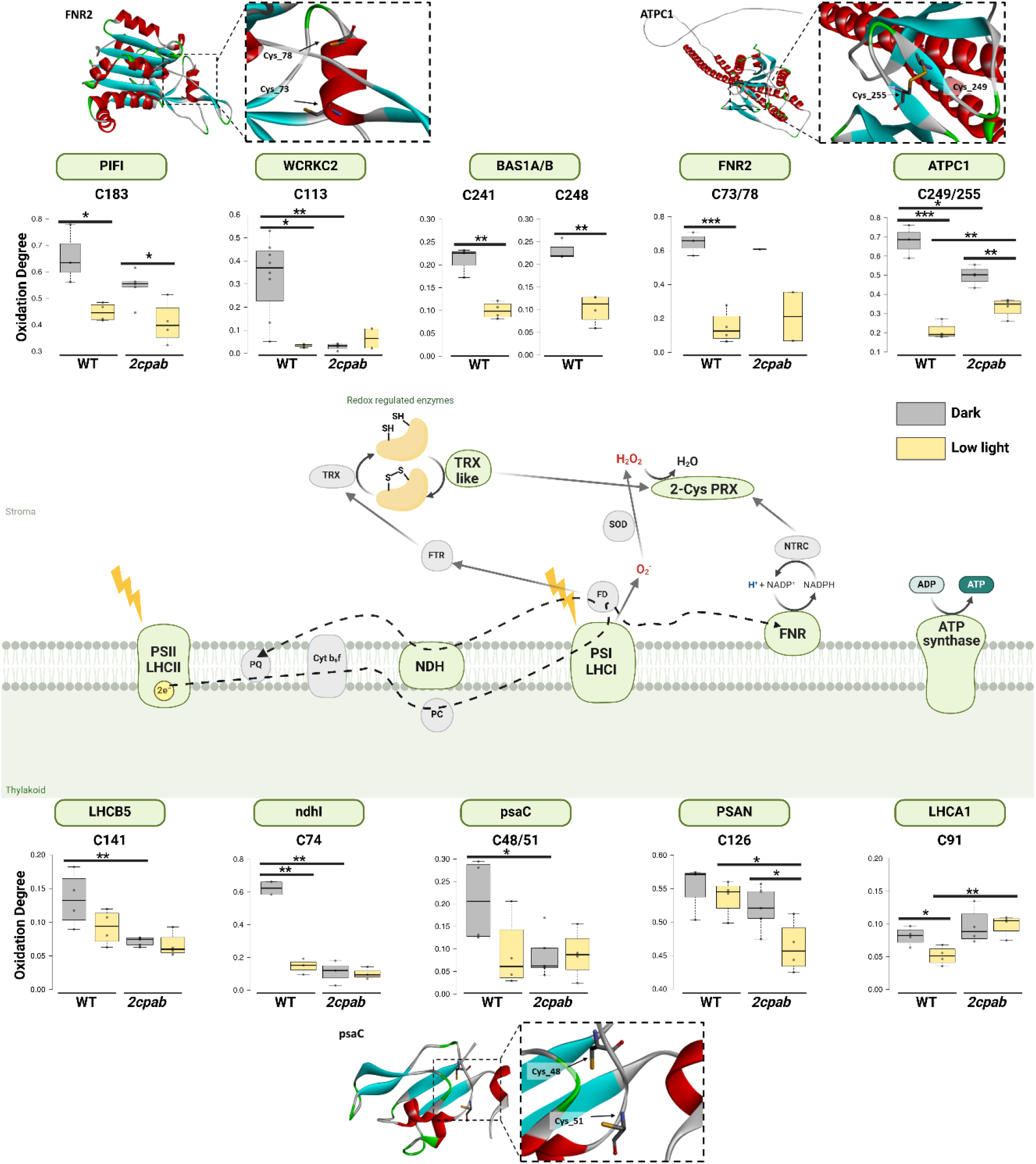
Redox-sensitive Cys sites in photosynthetic light reaction proteins. Schematic representation of the light-dependent reactions of photosynthesis. The oxidation degree of the specific identified Cys in dark and LL are presented as box plots for WT and *2cpab* plants. Statistical differences are stated through asterisks on the bars. For proteins: FNR2, ATPC1 and psaC, the oxidation degree was calculated for two identified Cys in the same protein, and the structure showing these Cys is presented.

Cysteines regulating the activity of CBC enzymes, including key regulatory sites in phosphoribulokinase (PRK) and glyceraldehyde-3-phosphate dehydrogenase subunit B (GAPB), were also reduced under LL in WT and were consistently reduced in *2cpab* (Fig.4). PRK and GAPB are redox-regulated through the formation and dissociation of a ternary complex that includes both enzymes and a conditionally disordered protein, CP12^40–42^. Specifically, Cys^243^ and Cys^249^ of PRK and Cys^434^ and Cys^443^ of GAPB, which participate in the formation of the GAPDH-CP12-PRK complex^43^, were reduced in response to LL in WT, but were highly reduced in the dark in *2cpab* plants and remained so in LL (Fig. 4). These results highlight the critical role of 2-Cys Prxs in regulating one of the key CBC regulation mechanisms under dark-light transition and in response to changes in light availability^42^ through the formation of GAPDH-CP12-PRK complex.

**Figure 4:**
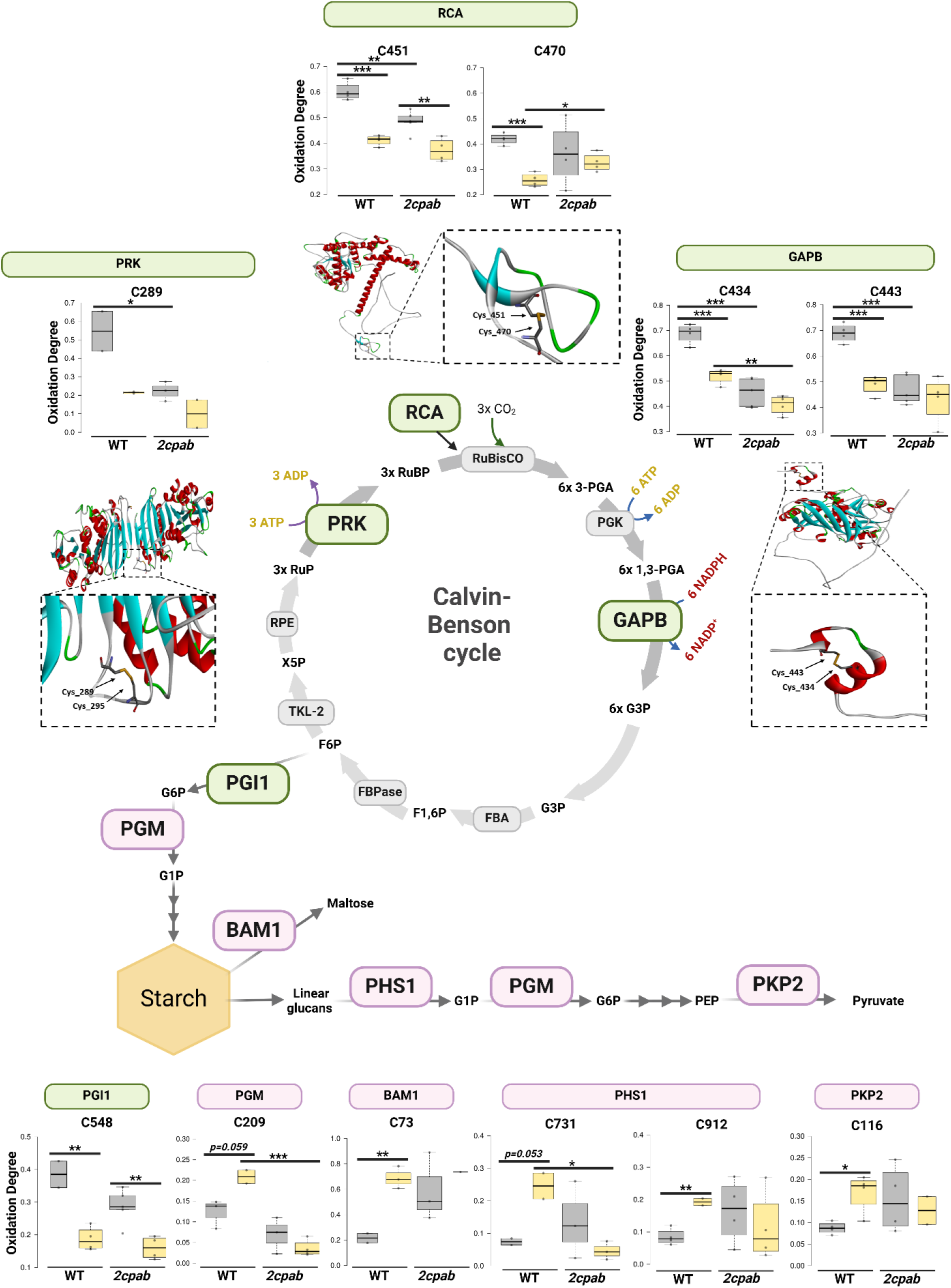
Redox-sensitive Cys sites in proteins involved in the Calvin-Benson cycle and downstream reactions. Schematic presentation of the Calvin-Benson cycle (CBC) and proteins involved in downstream reactions, i.e., starch synthesis, starch degradation and plastidial glycolysis. Cysteine sites that underwent reduction or oxidation during the transition from dark to LL in WT plants are colored green and pink, respectively. The oxidation degree of the identified Cys sites are presented in box plots for WT and *2cpab* plants under dark and LL conditions. Statistical differences are indicated by asterisks. For RCA, GAPB and PRK, the structures showing the participation of the identified Cys in forming disulfide bonds are presented.

An additional trend exhibiting Cys sites that were oxidized in the dark and reduced in response to LL in WT and *2cpab*, while exhibiting higher dark oxidation levels in WT compared to *2cpab* was identified. This suggests that dark-induced oxidation of these Cys residues involves both a 2-Cys Prx-dependent pathway and an alternative pathway that operates independently of 2-Cys Prx. These results align with the suggestion of a 2-Cys Prx-independent protein oxidation pathway^19,20,44^. Examples of cysteines in this group are Cys^249/255^ of ATP synthase gamma chain 1 (ATPC1), which are known to be redox-regulated and to form a reversible disulfide bridge^45^ as the one shown in Fig. 3. Another example is Cys^183^ of post-illumination chlorophyll fluorescence increase (PIFI) (Fig. 3).

Cysteines in two CBC-associated enzymes, ribulose bisphosphate carboxylase/oxygenase activase (RCA) and glucose-6-phosphate isomerase 1 (PGI1), also followed this trend. RCA is a key player in modulation of the overall carbon fixation level and the rate of induction of photosynthesis under changing light conditions^25^. Cys^451^ was reduced in response to LL in WT and *2cpab*, with the dark oxidation state being higher in WT than in *2cpab* (Fig. 4). The delayed oxidation of RCA in *2cpab* aligns with the role of 2-Cys Prxs in catalyzing its oxidation in the dark^20^. The OxD of Cys^470^, which forms the disulfide bond with Cys^451^, was found to be reduced in the transition to LL in WT. However, the effect of 2-Cys Prxs on its dark oxidation state was not statistically significant. PGI1 is a key player in the transitory starch biosynthesis in the leaf plastids^46^. The dark oxidation state of the redox-regulated Cys^548^ was found to be higher in WT compared to *2cpab*, with both plant lines exhibiting similar oxidation degrees under LL (Fig. 4).

### Light-induced oxidation of Cys regulatory sites in starch degradation and chloroplast glycolysis proteins

During the night, starch is degraded to sustain plant maintenance and growth in the absence of light. Similarly, chloroplast glycolysis is required to maintain metabolic reactions when photosynthesis does not occur^47^. Oxidation of several Cys sites in these pathways was observed in response to LL in WT plants. Cys^73^ in BAM1, a protein shown to be involved in starch degradation in guard and mesophyll cells, and whose activity is redox-regulated^48^, reached ΔOxD=50% in WT plants transferred to LL (Fig. 4). This cysteine forms a disulfide bond with Cys^511^ (not identified in our data), and is inhibited under oxidizing conditions^46^. Similar redox behavior was observed in Cys^731^ and Cys^912^ of α-glucan phosphorylase 1 (PHS1), a starch-degrading enzyme involved in the phosphorolytic pathway^46^ (Fig. 4). Light-induced oxidation of Cys^116^ of plastidial pyruvate kinase 2 (PKP2), which catalyzes the irreversible synthesis of pyruvate and ATP in plastidial glycolysis, was also detected. Similarly, Cys^209^ of putative phosphoglucomutase (PGM), which catalyzes the reversible conversion of glucose-1-phosphate (G1P) to glucose-6-phosphate (G6P), playing a role in metabolism of carbohydrates derived from starch degradation as well as in starch biosynthesis, underwent oxidation in WT under LL. No light-induced oxidation of these Cys sites was observed in *2cpab* plants, with some sites, such as Cys^209^ in PGM and Cys^731^ in PHS1, remaining reduced in both dark and LL, while others exhibited a highly oxidized state under both conditions. Taken together, these data indicate light-induced oxidation of Cys sites in enzymes involved in carbohydrate catabolism and suggest that the redox regulation of these pathways is disrupted in *2cpab* plants.

## Discussion

This work aimed to systematically map redox changes at the proteome level during photosynthesis induction and to unravel the role of 2-Cys Prxs in shaping these redox alterations. The analysis provided three key insights into redox events occurring during the induction phase: 1) Redox modifications during the induction phase include both reductive and oxidative signals, sensed by distinct sets of proteins. 2) Oxidative siganls are regulated by 2-Cys Prx, while an additional 2-Cys Prx-independent mechanism is involved in protein oxidation of certain Cys sites. 3) The availability of substrates for reductive signals depends on 2-Cys Prx oxidative activity in the dark.

Chloroplast metabolism must be tightly regulated during transitions from dark to light and shifts from low to high light. In the light, enzymes driving light and carbon reactions and starch biosynthesis are activated, while those involved in starch degradation and possibly chloroplast glycolysis are deactivated. The opposite response occurs when the light is turned off: the day metabolic proteins are deactivated, while nocturnal metabolic pathways are reactivated. A mechanism possibly enabling this redox-response specificity involves a temporal separation between opposing redox signals, with reductive activity by the Fd/FTR pathway occurring in the light and oxidative activity by atypical TRXs/2-Cys Prxs occurring in the dark. In this manner, specificity is determined by the inherent properties of target proteins, specifically, the correlation between their redox state and their functional efficiency. Accordingly, photosynthetic enzymes are activated in their reduced states when light is available, while enzymes involved in dark metabolism are activated by oxidation, which predominates when the plant is not actively photosynthesizing. Indeed, deactivation upon reduction and activation upon oxidation of glucose 6-phosphate dehydrogenase and phosphofructokinase 5 (PFK5), which are involved in the pentose phosphate and chloroplast glycolytic pathways, respectively, has been demonstrated^49,50^. However, temporal separation of reductive and oxidative activities, along with the unique characteristics of target proteins, does not fully explain specificity, as only a few proteins were found to be inactivated upon reduction, and counterintuitively, many starch-degradation enzymes required in the dark are activated by reduction^51^, suggesting that protein properties alone cannot account for this regulation. Indeed, several observations indicate that reductive and oxidative signals arise concurrently. For example, the simultaneous generation of ETR-related reductive and oxidative signals during the dark-to-light transition was demonstrated using genetically encoded redox-sensitive biosensors^31,52^. In addition, protein oxidation measurements have shown that despite the expected reduction of many target proteins, several components of the oxidative pathways, such as ACHT1/4 and 2-Cys Prx, as well as with NTRC, and Trx-y2, which function as electron donors for 2-Cys Prx^17,18,53^, are oxidized upon light exposure. A rise in oxidation state upon illumination was also demonstrated for chloroplast superoxide dismutase [Cu-Zn] 2 (CSD2) and several proteins residing outside the chloroplast^29,53^. These observations are consistent with the presented data, which show the simultaneous reduction and oxidation of Cys residues in different enzymes.

The emerging intricate and finely tuned redox mechanism in which opposing signals are generated simultaneously and affect different sets or regulatory sites implies the existence of hubs, possibly membrane-less organelles, within the stromal environment, that confer specificity in the reception of distinct redox signals. Such partitioning of the chloroplast environment into redox-specific nanoscale regions may also explain the unexpected transmission and reception of oxidative signals during the induction phase, when light absorption at photosystems stimulates low-redox-potential environments. Similarly, protein-context-dependent redox differences, demonstrating unique “redox microdomains”, were recently suggested, based on highly specific and dynamic changes in H_2_O_2_ availability on the scale of individual protein complexes, observed using a yeast library in which the H_2_O_2_ biosensor was fused to all protein-coding open reading frames^54^. The specificity of different redox hubs may be governed by the selective engagement and release of specific Trxs, with redox outcomes determined by the redox potentials and binding specificity of the Trx. For example, a hub integrating an atypical Trx, while excluding Trxs that drive reductive reactions, may create a local environment that favors oxidative activity.

Light-induced reduction and oxidation were disrupted in *2cpab* plants, pointing to the role of 2-Cys Prxs in shaping the redox proteome during the induction phase. While 2-Cys Prx directly facilitates protein oxidation in the light, it indirectly modulates reductive signals by facilitating substrates for reduction through its oxidizing activity in the dark. The fact that many photosynthetic regulatory cysteines remained fully or partially reduced in the dark may explain the rapid induction of photosynthesis in *2cpab* plants^31^, with the continuously active state of light and carbon reactions facilitating the swift activation of CO_2_ metabolism, shortening the time required to activate photosynthetic proteins. However, the reduced growth and lower chlorophyll content observed in *2cpab* plants compared to WT^14^ point to the critical role of oxidative activity in cellular homeostasis and suggest that maintaining metabolic proteins in a continuously “on” state comes with a metabolic price. Specifically, the disrupted redox regulation of starch degradation and chloroplast glycolysis enzymes, such as BAM1 and PKP2, suggest impaired energy and carbon flux regulation, which may result in energy waste and subsequently slower growth rates. Indeed, synchronization of light absorption in reaction centers, carbon reactions and the starch biosynthesis/degradation cycle, has been shown to be essential for optimizing plant productivity^55^. The possibly impaired regulation of cyclic electron flow in *2cpab*, as reflected by the highly reduced state of Cys^74^ in NdhI, may also account for the stunted growth of the mutant (Fig. 3). Further biochemical investigations are required to precisely characterize the metabolic costs associated with disrupted redox regulation in *2capb* plants.

In their natural environment, plants are exposed to dynamic and fluctuating light conditions, necessitating continuous finetuning of metabolic fluxes to maintain homeostasis. Consequently, precise generation and perception of reductive and oxidative signals are essential, not only during dark-light transitions but also when adapting to varying light intensities. Given the pivotal role of redox regulation in modulating photosynthetic activity under such conditions, elucidating the mechanisms that confer specificity may provide a foundation for future advancements in optimizing the photosynthetic process.

## Materials and Methods

### Plant Material, growth conditions and treatment

*Arabidopsis thaliana* WT plants (ecotype Columbia, Col-0) were obtained from ABRC. The mutant *2cpab* was obtained from Francisco Javier Cejudo, Universidad de Sevilla, Sevilla, Spain. WT and *2cpab* mutant plants were grown in a greenhouse under a 16/8 h light/dark cycle. Immunoassay and thiol labeling experiments were performed on 3-week-old whole Arabidopsis plants.

### Immunoassay

Protein extract was prepared in 2 ml tubes using a hand homogenizer in the presence of 1 ml cold 10% trichloroacetic acid (TCA) dissolved in water. Proteins were precipitated for 30 min on ice in the dark and centrifuged at 14,000 rpm, for 20 min, at 4°C. The pellet was then washed 4 times with 100% cold acetone. Residual acetone was removed and the pellet was resuspended in urea buffer comprised of 8 M urea, 100 mM 4-(2-hydroxyethyl)-1-piperazineethanesulfonic acid (HEPES) (pH 7.2), 1 mM EDTA, 2% (w/v) SDS, protease inhibitors cocktail (PI) (Calbiochem) and 100 mM N-ethylmaleimide (NEM) (E3876, Sigma) dissolved in ethanol. The samples were incubated for 30 min at room temperature and then centrifuged (14,000 rpm, 20 min, 4°C) and washed 4 times with 100% cold acetone. The dry pellets were resuspended in urea buffer without NEM and then fractionated on a 4–15% precast polyacrylamide gel without reducing agent. Sample buffer (x3) was comprised of 150 mM Tris-HCl, pH 6.8, 6% (w/v) SDS, 30% glycerol and 0.3% pyronin Y. Fractionated proteins were transferred onto polyvinylidene fluoride membrane (Bio-Rad), using a Trans-Blot Turbo Transfer System (Bio-Rad) with Trans-Blot Turbo Midi Transfer Packs. The membrane was incubated with anti-Prx antibodies (1:1,000) (was kindly provided by Prof. Avichai Danon), followed by secondary anti-rabbit horseradish peroxidase (1:20,000) (Agrisera), and later developed using standard protocols.

### Thiol-labeling assay

Protein extract was prepared in 2 ml tubes using a hand homogenizer, in the presence of 1 ml cold 20% TCA dissolved in water. Proteins were precipitated for 30 min on ice in the dark, and centrifuged at 14,000 rpm for 20 min, at 4°C. The pellet was washed once with 10% TCA (dissolved in water) and 2 times with 100% cold acetone. The residual acetone was removed by brief evaporation and the precipitate was resuspended (600 µl) in urea buffer containing 100 mM N-ethylmaleimide (d_0_-NEM) (E3876, Sigma), dissolved in ethanol, to label the reduced cysteines. Samples were incubated at 50°C for 30 min, with 1000 rpm shaking. The reaction was terminated, and proteins were precipitated by bringing the TCA content to 20% (30 min on ice in the dark), washed once with 10% TCA and twice with 100% acetone. Samples were then resuspended in urea buffer supplemented with 5.5 mM tris 2-carboxyethyl phosphine (TCEP, Sigma) but without NEM, and shaken (1200 rpm, 50°C, 30 min). Samples were further alkylated by adding 5 mM (final concentration) d5-NEM (DLM-6711, CIL) and incubated (50°C, 30 min, 1200 rpm shaking). TCA (33%) was added, samples were precipitated for 30 min on ice in the dark and then centrifuged (14,000 g, 4°C, 20 min). The pellet was then washed once with 10% TCA and twice with 100% acetone. The acetone was removed by 5 min evaporation and samples were stored at -80°C.

Four biological replicates were performed for dark and low light (LL) treatments each, for WT and *2cpab*, resulting in sixteen-core samples. Two separate experiments were conducted to increase the number of identifications, which amounts to 160 LC-MS/MS runs. The final peptide list included unique peptides from each experiment, and in cases where a peptide was identified in both experiments and quantification of the oxidation degree was available in at least two replicates per condition/plant line, the experiment exhibiting smaller *p-value* in the ΔOxidation degree (LL-dark) of the WT plants was selected. The data of all replicates collected from the two experiments is available as Supp.Table 3.

An additional set of four samples for each experiment (eight samples in total for both experiments) were subjected to thiol labeling as described above, however, unlike the former labeling, the reduced and oxidized cysteines were all labeled with d_5_-NEM. Following precipitation with 20% TCA, the samples were resuspended (250 µl) in urea buffer, reduced with 5.5 mM TCEP and further alkylated by adding 100 mM d5-NEM (dissolved in ethanol) (final concentration). The samples were stored at -80°C. Regarding the LC-MS/MS, 40 runs were added to the number mentioned above.

### Sample preparation for LC-MS/MS analysis

All chemicals were from Sigma Aldrich, unless otherwise indicated. Alkylated protein pellets were dissolved by addition of lysis buffer (5% SDS, 50 mM Tris-HCl, pH 7.4), heating for 15 min at 50°C, sonication, and incubation at room temp for 4 h. Alkylated proteins were suspended in lysis buffer (5% SDS, 50 mM Tris-HCl, pH 7.4). After determination of protein concentration using the BCA assay (Thermo Scientific, USA), 100 μg total protein of each sample was loaded onto S-Trap mini-columns (Protifi, USA), according to the manufacturer’s instructions. In brief, after loading, samples were washed with 90:10% methanol/50 mM ammonium bicarbonate and then digested (1.5 h, 47 °C) with trypsin (1:50 trypsin/protein). The digested peptides were eluted using 50 mM ammonium bicarbonate and then incubated with trypsin overnight at 37°C. Two more elution rounds were performed using 0.2% formic acid and 0.2% formic acid in 50% acetonitrile. The three eluted fractions were pooled and vacuum-centrifuged to dryness. Samples were stored at −80 °C until analysis.

### Liquid chromatography

ULC/MS-grade solvents were used for all chromatography steps. Each sample was fractionated offline using high-pH reversed-phase separation, followed by online low-pH reversed-phase separation. Digested protein (100 µg) was loaded using high-performance liquid chromatography (H-Class, Waters). Mobile phase was: A) 20 mM ammonium formate pH 10.0, B) acetonitrile. Peptides were separated on an C18 XBridge column (3.0×150 mm, Waters). Column temperature was set to 45°C, flow rate of 0.3 mL/min and the following gradient was used: 3% B for 2 min, linear gradient to 40% B over 50 min, maintained at 40% B for 5 min, 5 min to 95% B, and then back to the initial conditions. Peptides were fractionated into 14 fractions, each of 1.2 mL. The fractions were dried by speed-vac and reconstituted in 40 µL solution of 97% water, 3% acetonitrile, 0.1% formic acid. The following fractions were then pooled: Fr1: 1,2,3,8; Fr2: 4,9; Fr3: 5,10,11; Fr4: 6,12,13,14; Fr5:7. Each fraction was dried in a speedvac, then reconstituted in 40 µL solution of 97% water, 3% acetonitrile, 0.1% formic acid. Each pooled fraction was then loaded and analyzed using split-less nano-ultra performance liquid chromatography (10 kpsi nanoAcquity; Waters, Milford, MA, USA). The mobile phase was: A) H_2_O + 0.1% formic acid and B) acetonitrile + 0.1% formic acid. Samples were desalted online using a Symmetry C18 reversed-phase trapping column (180 µm internal diameter, 20 mm length, 5 µm particle size; Waters). The peptides were then separated using a T3 HSS nano-column (75 µm internal diameter, 250 mm length, 1.8 µm particle size; Waters) at 0.35 µL/min. Peptides were eluted from the column into the mass spectrometer, using the following gradient: 4% to 27% B over 105 min, 27% to 90%B over 5 min, maintenance at 90% for 5 min and then back to the initial conditions.

### Mass spectrometry

The nanoUPLC was coupled online through a nanoESI emitter (10 μm tip; New Objective; Woburn, MA, USA) to a quadrupole orbitrap mass spectrometer (Orbitrap Fusion Lumos, Thermo Scientific) using a FlexIon nanospray apparatus (Proxeon). Data were acquired in DDA mode, using a 2-sec cycle method. MS1 was performed in the Orbitrap with resolution set to 120,000 (at 200 m/z) and maximum injection time set to 246 msec. MS2 was performed in the Orbitrap after HCD fragmentation, with resolution set to 15,000 and maximum injection time set to 60 msec.

### Data analysis -protein expression

Raw data was processed with the MetaMorpheus v1.0.2 informatics platform. The data were searched against the Arabidopsis proteome database as downloaded from Uniprot.org, and appended with common lab protein contaminants. Variable modifications were defined to include N-ethylmaleimide on C and Nem:2H(5) on C, as well as N-ethylmaleimide+ water on C and Nem:2H(5)+H_2_O on C. Peptide intensities are reported by MetaMorpheus for each fraction separately. To generate the final peptide table, the intensities from all 5 fractions were summed up for each sample. The quantitative comparisons were calculated using Perseus v1.6.15.0. Decoy hits were filtered out, and only proteins/peptides that were identified in at least 50% of the replicates of at least one experimental group were kept.

### Data analysis – Cys oxidation

Peptide identification and quantification were conducted using the MetaMorpheus software (v0.0.297 https://smith-chem-wisc.github.io/MetaMorpheus/ or https://github.com/smith-chem-wisc/MetaMorpheus). RAW MS files were searched against the *Arabidopsis thaliana* (Mouse-ear cress) proteome downloaded from UniProt (UniProtKb Proteome ID UP000006548, as of June 13, 2023, in XML format). All MS files, including the sample in which reduced thiols were labeled with d0-NEM and those labeled with d_5_-NEM, were jointly processed. In short, the MetaMorpheus calibration task was performed using the default parameters. In the search task, we used N-ethylmaleimide on C and Nem:2H(5) on C, as well as N-ethylmaleimide+ water on C and Nem:2H(5)+H_2_O on C as variable modifications. Moreover, the following options were checked in the search settings: apply protein parsimony and construct protein groups, treat modified peptides as different peptides, match between runs, and normalize quantification results.

The generated output files ‘AllQuantifiedPeptides.tsv’ were used for further analysis. First, files were trimmed, and only cysteine-containing peptides labeled with d_0_ or d_5_-NEM, as well as d_0_ or d_5_-NEM + water (belongs to the core set in which reduced Cys was labeled with d_0_-NEM) were selected. Next, a list with peptides bearing both labels was extracted (pairs). A CysID was then assigned to each cysteine-containing peptide; the ID included the UniProt identifier name of the protein and the NEM-labeled cysteine residue position in the protein sequence. Next, the intensity of each peptide throughout the five fractions was summed. If a specific Cys was represented by more than one peptide, the overall intensity of peptides with identical CysIDs was examined and the peptide with the highest intensity was selected for further analysis. The oxidation degree (OxD) of a Cys was calculated using the following formula: OxD=d_5_/(d_5_+d_0_), where d_5_ represents the intensity of the d_5_-NEM labeled peptide and d_0_ the intensity of the d_0_-NEM labeled peptide. Only peptides identified in at least two out of four biological replicates were included. Delta oxidation (ΔOxD) was calculated by subtracting the OxD of LL-treated plants from the OxD of dark-treated plants. The p-value for each peptide was determined using student’s t-test.

The OxD for peptides with 2 Cys was calculated using the formula:

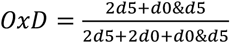, where 2d5 represents the intensity of the peptide with 2 Cys that both Cys were d_5_-NEM labeled, 2d_0_ the intensity of the peptide with 2 Cys that both Cys were d_0_-NEM labeled, and d_0_&d_5_ the intensity of the peptide with 2 Cys that one Cys was d_0_-NEM labeled and the other Cys was d_5_-NEM labeled. The two corresponding ΔOxD values were calculated and the p-value for each peptide was determined using student’s t-test, accordingly.

Only peptides that passed the threshold of ΔOxD > |±5%| and α<0.05 were considered “redox-sensitive peptides”.

### Functional annotation analysis

The DAVID (Database for Annotation, Visualization and Integrated Discovery) Bioinformatics Resources tool v6.7 was used for mapping protein into Gene Ontology terms (GO) and KEGG (Kyoto Encyclopedia of Genes and Genomes) pathways and for enrichment analysis. Further mapping into metabolic pathways was carried out using the Arabidopsis chloroplast knowledge base (cloroKB, ^56^), the AraCyc database from the Plant Metabolic Network (PMN^57^), MapMan, and manual annotation. Subcellular protein localization was predicted using the SUBA tool (the central resource for Arabidopsis protein subcellular location data). Venn diagrams depicting the associations and distinctions of the reduced and oxidized unique cysteine-containing peptides to the transition from dark to LL in the WT vs *2cpab* Arabidopsis mutant plants were generated using the Venny 2.1 tool (https://bioinfogp.cnb.csic.es/tools/venny/). ΔOxD of Cys belonging to various subcellular compartments to the same cluster that show the same phenomenon was presented using the PlotsOfData tool (https://huygens.science.uva.nl/PlotsOfData/). Oxidation degree of specific Cys of interest, and ΔOxD of Cys belonging to the same biochemical pathway were presented using the BoxPlotR: a web-tool for generation of box plots tool (http://shiny.chemgrid.org/boxplotr/). The Probability logo generator for biological sequence motifs (pLogo, https://plogo.uconn.edu/^58^) was used to estimate the probability of the occurrence of specific amino acid sequence motifs around the NEM labeled Cys. The functional annotation regarding the active and binding sites, metal binding and disulfide bonds, for all the quantified cysteines was retrieved from the UniProt Knowledgebase (https://www.uniprot.org/uploadlists/). Discovery Studio 4.5 Visualizer was used for structure visualization of Cys location in protein structures. All images were created with BioRender (https://biorender.com/).

## Supporting information

Supplementary Images 1-3

