## Supplementary Images 1-3 for "Two opposing redox signals mediated by 2-Cys peroxiredoxin shape the redox proteome during photosynthetic induction"

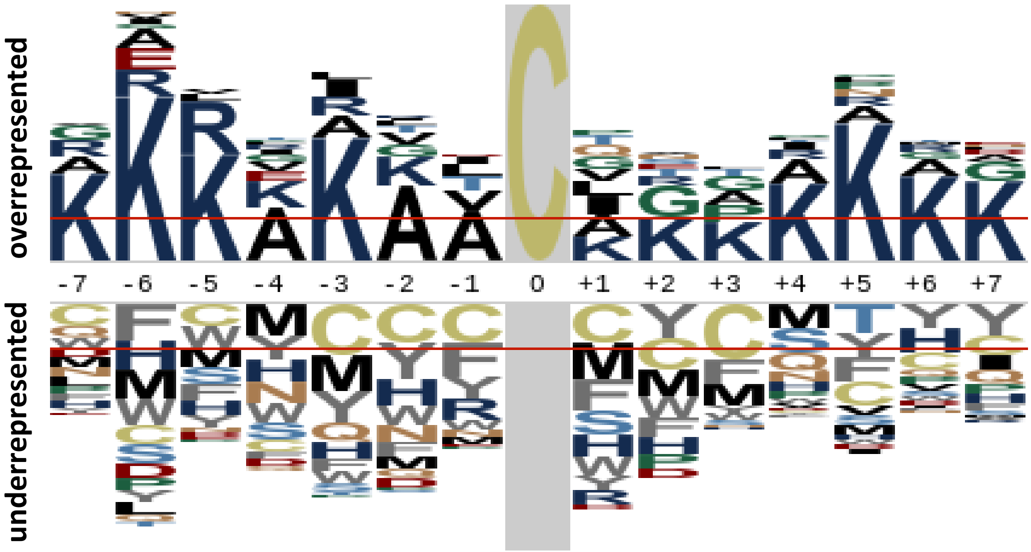


**Supplementary Figure 1: Consensus motif of identified Cys containing peptides not labeled with NEM.** *Arabidopsis thaliana* proteome was used as the proteomic background population for the calculation of the probabilities.

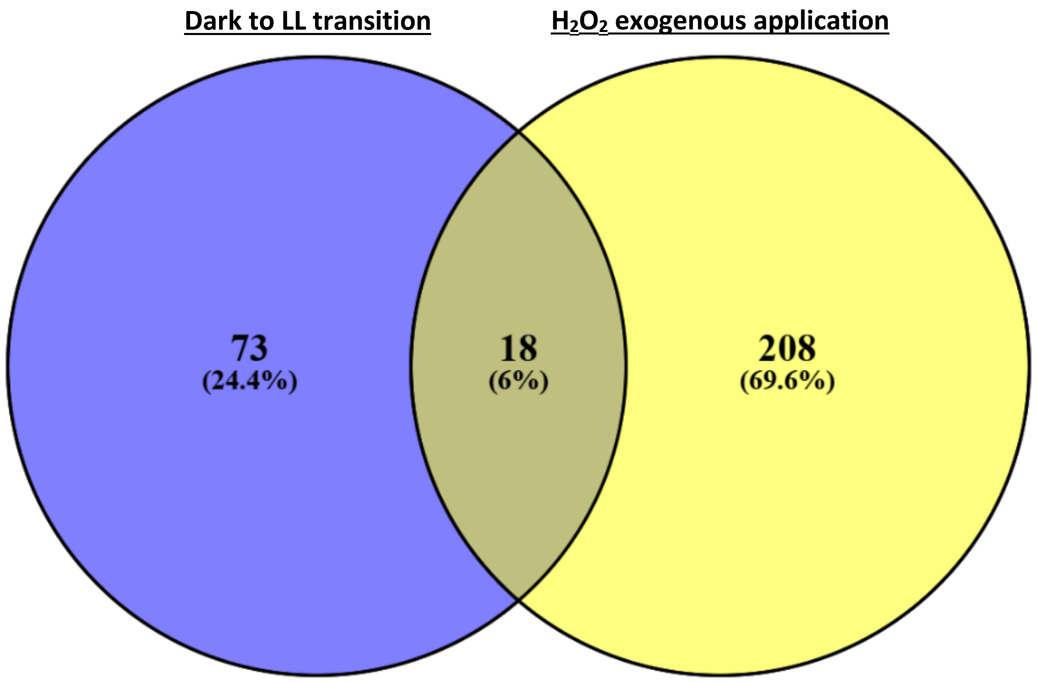


**Supplementary Figure 2.** Venn diagram illustrating the overlap between redox-regulated cysteine-containing peptides in WT plants during the transition from dark to low light (LL) (this study) and in response to exogenous $H_2O_2$ application (Doron et al., 2021). Only peptides identified in both experiments were included.

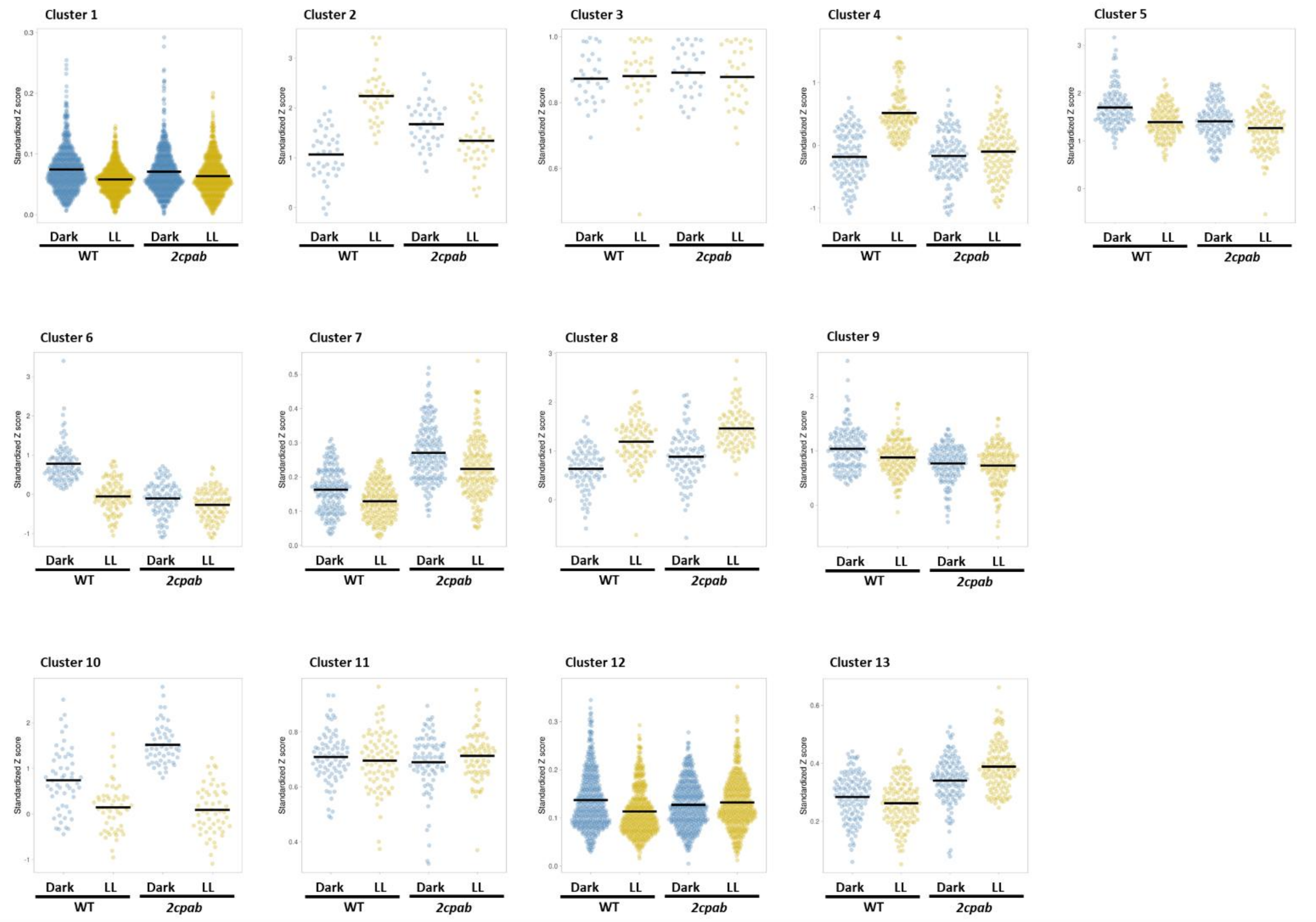


**Supplementary Figure 3**. Violin plots representing oxidation degree levels across the clusters shown in Fig. 2c. Each plot corresponds to a distinct cluster, illustrating the distribution of Cys oxidation values within that cluster.
